# Olfactory search with finite-state controllers

**DOI:** 10.1101/2023.03.10.532037

**Authors:** Kyrell Vann Verano, Emanuele Panizon, Antonio Celani

## Abstract

Long-range olfactory search is an extremely difficult task in view of the sparsity of odor signals that are available to the searcher and the complex encoding of the information about the source location. Current algorithmic approaches typically require a continuous memory space, sometimes of large dimensionality, which may hamper their optimization and often obscure their interpretation. Here, we show how finite-state controllers with a small set of discrete memory states are expressive enough to display rich, time-extended behavioral modules that resemble the ones observed in living organisms. Finite-state controllers optimized for olfactory search have an immediate interpretation in terms of approximate clocks and coarse-grained spatial maps, suggesting connections with neural models of search behavior.

## 1. Introduction

Many living organisms rely on odor signals to locate distant sources of nutriment or to approach potential mates [1, 2, 3, 4, 5, 6]. Their search strategies crucially depend on how the information is transmitted across large distances by the atmospheric flow. Turbulent transport leads to a dynamic, complex and sparse odor landscape far away from the source, with time intervals between consecutive odor encounters that go from milliseconds to minutes [7]. The information about the source location is somehow encoded in this complex sequence of odor concentrations detected by the searcher [8, 9]. The navigation strategies adopted by insects in these situations are quite structured. For instance, in moths, the search patterns are often characterized by extended crosswind movements (“cast”) alternated to stretches of upwind flight (“surge”), as sketched in Figure 1. It has been experimentally observed that the movement of the moth is strongly influenced by the frequency of past detections: repeated odor encounters lead to longer surges and sparser ones are conducive to wider casts [10, 11].These observations point to the conclusion that the navigation strategy depends in a nontrivial way from the history of past odor encounters and therefore requires some form of memory.

**Figure 1:**
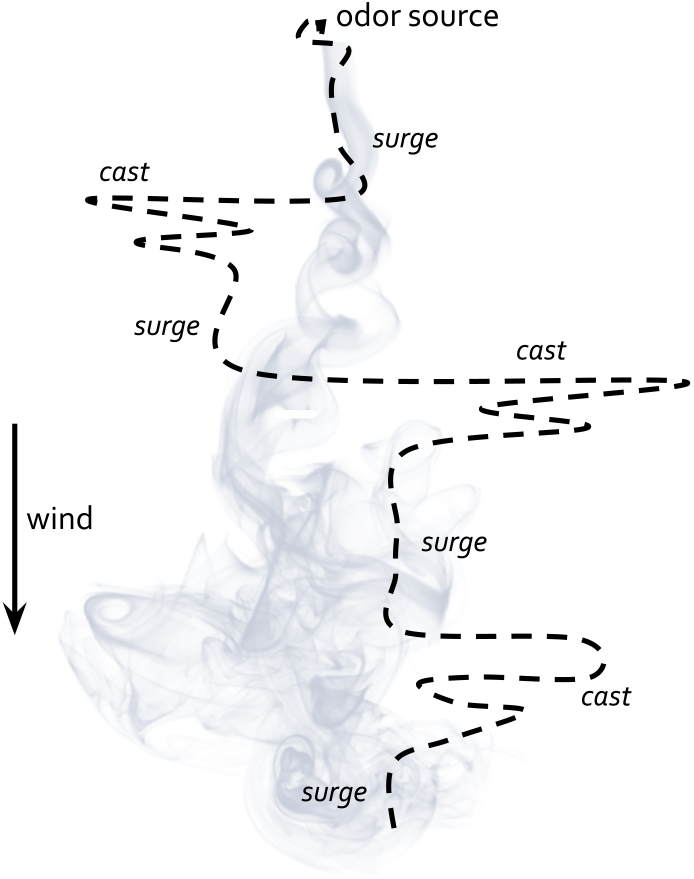
A sketch of a trajectory of a moth searching for an odor source in a turbulent plume.

In this paper we study the structure of memory in algorithmic models of olfactory search. We identify minimal models, that take the form of *finite-state controllers* (FSC), which have a discrete memory space and can be efficiently optimized. We find that three or four memory states are already expressive enough to account for elaborate navigation patterns that resemble the strategies observed in living organisms – or the ones produced by much more complex algorithms. In the FSC optimized for olfactory search that we show below, the memory states can be straightforwardly interpreted as approximate clocks and coarse-grained spatial maps, suggesting a possible connection with the neural processes at work in the brain of insects and other organisms.

### 1.1. The algorithmic view of olfactory search

At the most general level, olfactory search is a specific instance of a Partially Observable Markov Decision Process (POMDP) [12, 13]. The structure of a POMDP is illustrated in Fig. 2. The world is partitioned in three compartments: the agent, the environment, and the sensorimotor interface. The agent is the decision-maker: we call its internal state the *memory* and label it by the symbol m. The interaction of the agent with the outer environment is mediated by the sensorimotor interface. Information about the *state s* of the external world is collected by means of sensors that receive *observations y*. The agent can make decisions – on the basis of its current state and the observation that it has just received – in the form of actions a. At the same time, the memory is updated by incorporating the observation that has just been received, and as a result of the action, the environment can change its state. As is clear from the diagram, the process that evolves (*s_t_, m_t_*) into (*s*_*t*+1_, *m*_*t*+1_) has the Markov property.

**Figure 2:**
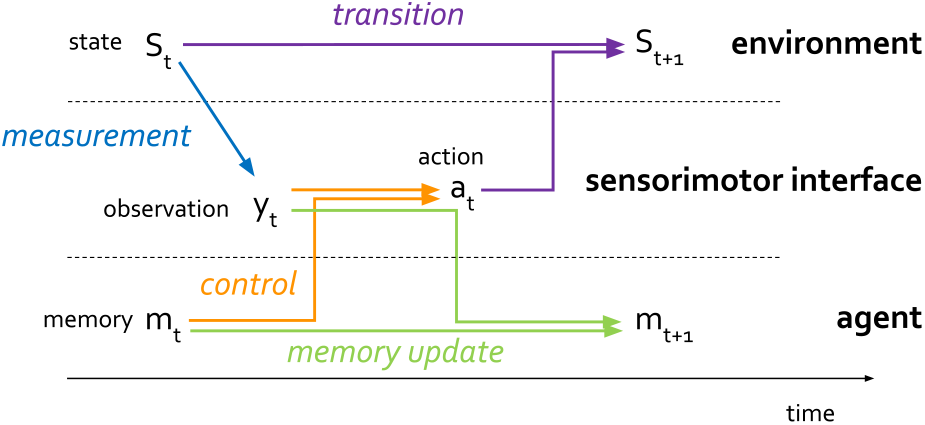
General scheme of a POMDP

The final and defining ingredient of a POMDP is the goal: the agent aims at maximizing, in expectation, the cumulative sum of *rewards r* that it can collect over time. Rewards are (possibly random) functions of the current state of the environment, of the action, of the successive state of the environment and of the observation: *r*(*s_t_, a_t_, s*_*t*+1_, *y_t_*).

This is a very general description that encompasses most, if not all, decision processes. For olfactory search we have the following correspondences:

- The agent is an abstraction of a living organism or an autonomous vehicle equipped with sensors and actuators.
- The state of the environment is in general a very highdimensional vector that contains, in addition to the position of the source, the location, pose and speed of the searcher, the concentration of odor, the velocity of the fluid at all points in space at a given time, and possibly more. The knowledge of all these variables is necessary to determine the state at the subsequent time. In other words, they are a Markov state for the dynamics of the environment conditioned on actions.
- Observations only give a very limited amount of information about the full state of the environment. Typically they include noisy measurements of odor concentration and of the wind direction together with some visual cue enabling the agent to assess its relative speed with respect to the ground.
- Actions are limited by the manoeuvring ability of the searcher. In most cases they include the possibility of making a turn and change speed.
- Rewards are often defined in terms of minus the time elapsed during one step of the search process, so that the overall goal is to minimize the average time needed to reach the source of odor.
- The memory space 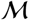 is in general designed with the ambition of emulating the capabilities of a brain. In practice, however, it can be as a discrete set, a subset of a vector space or a function space.

Since the structure of the memory space is the key subject of this work, in the next subsection we will provide detailed explicit examples for the case of olfactory search.

### 1.2. The uses of memory for olfactory search

We will focus below on three different ways of constructing a POMDP, each making use of a very different memory space and its corresponding memory updates. We will compare these different methods in the context of a simplified model of olfactory search on a two-dimensional rectangular grid, where the searcher can move along the principal directions and observations are binary, corresponding to the detection of an odor, or the lack thereof. Although this is obviously a very crude approximation of the actual process taking place in the real world, this model is customarily because it helps in highlighting the algorithmic aspects of the problem. In Table 1 we provide a precise characterization of the environment along with a glossary of the various definitions used in the paper.

**Table 1:**
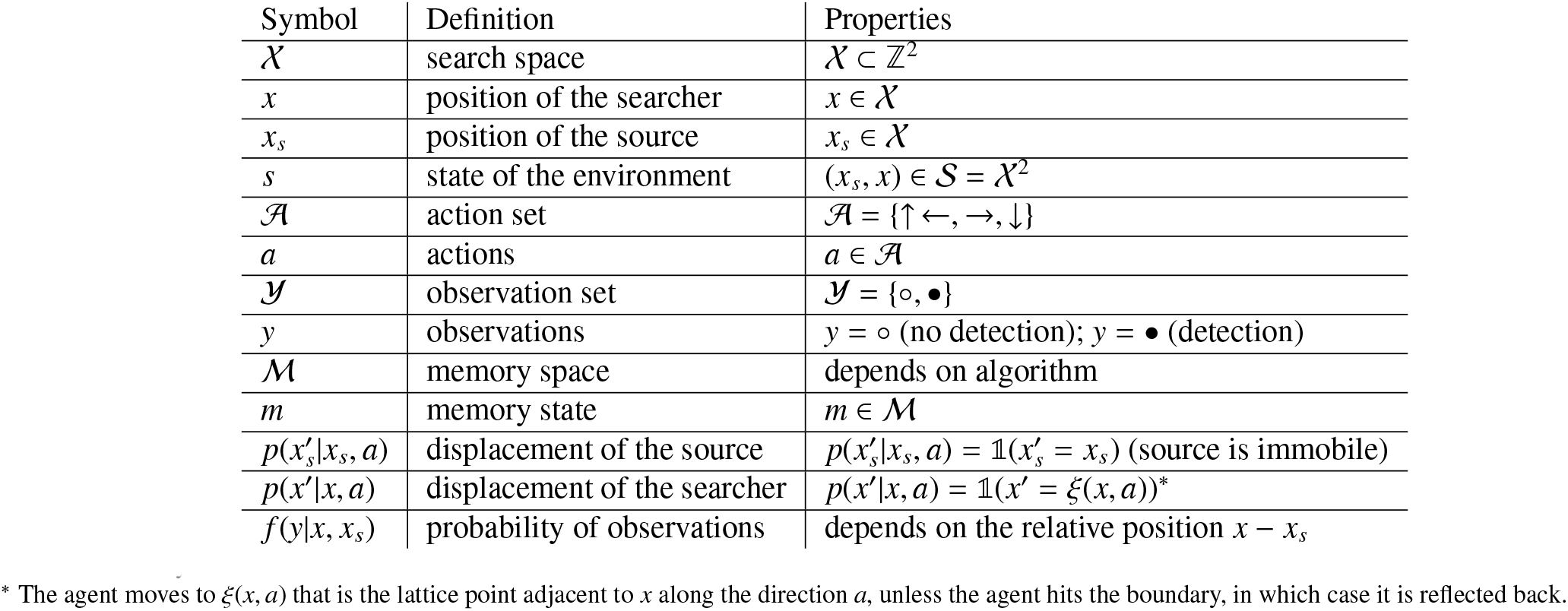
Glossary and definitions. * The agent moves to ξ(*x, a*) that is the lattice point adjacent to *x* along the direction *a*, unless the agent hits the boundary, in which case it is reflected back.

#### 1.2.1. The “Cast-and-Surge” algorithm: memory as a clock

We start with a heuristic algorithm that is an abstraction of the phenomenology of olfactory search depicted in Fig. 1 and was first introduced in Ref [10]. Here we rephrase it in the POMDP language to clarify its relationship with other approaches.

In this setting, the memory space is discrete and consists of all integer numbers: 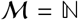. The memory updates and action choices are deterministic and the process in memory space is represented by the graph in Fig. 3. When there is an odor encounter (*y* = •), the memory is updated to *m* = 0 (dashed green arrows) and the agent moves upwind (*a* =↑). Conversely, when the odor is not detected, the memory is updated according to *m* → *m* + 1 (full green arrows) and the corresponding action is taken (orange arrows). Following this algorithm, the agent moves upwind when it detects the odor and enters a program made of alternating casts of increasing duration, according to a predetermined rule. Clearly, in this case the memory acts as a clock that is reset every time the agent detects the signal.

**Figure 3:**
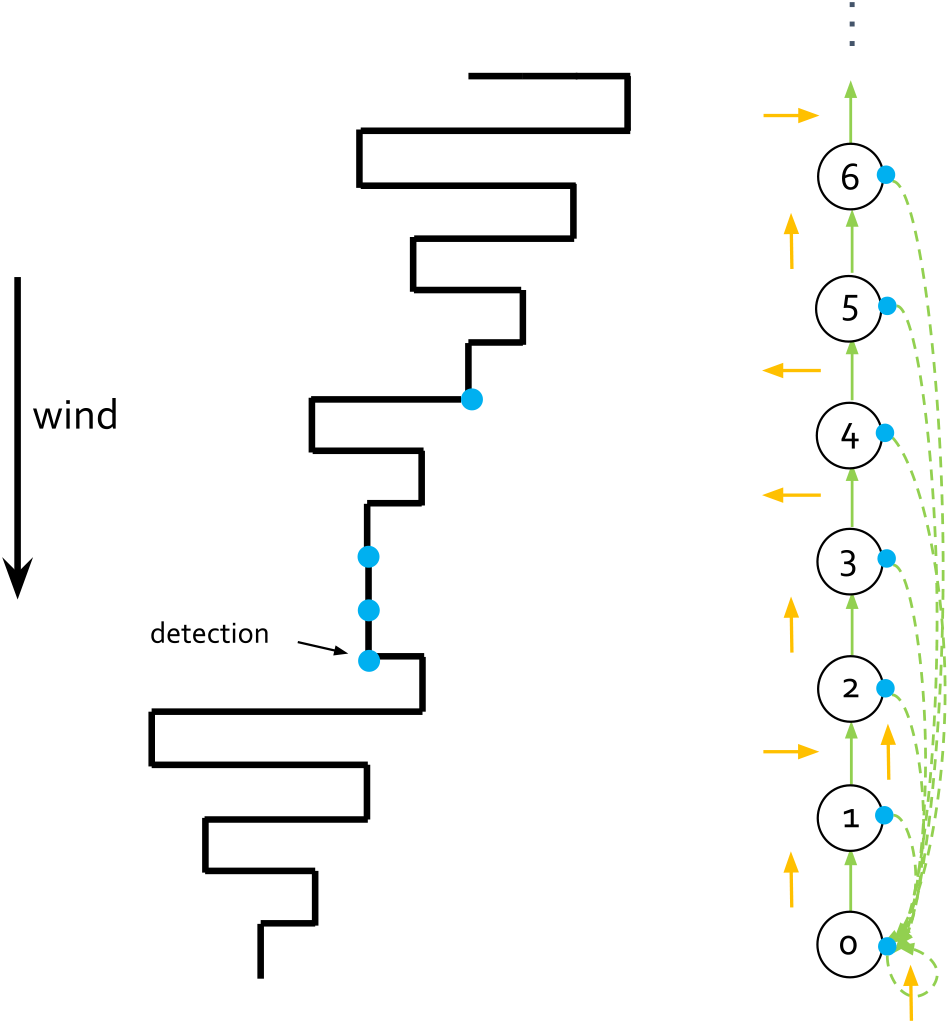
Graphical scheme of the “cast-and-surge” algorithm. On the left, a typical trajectory with surges and increasingly wide horizontal casts. Blue dots denote detection events. On the right we show the process in memory space.

It is important to remark that this strategy is *hardwired*, in the sense that there is no (or very limited) attempt at the optimization of performance. It is also *model-free* in that its implementation does not make use of detailed knowledge of the environment such as the space-dependence of the probability of detection.

#### 1.2.2. Bayesian search: memory as a map

In the Bayesian approach to POMDP, the unknown state of the environment is treated as a random variable with a probability distribution b(s) that is called the *belief* [12, 14, 15]. The memory state is the belief itself *b* and the memory space is the space of probabilities over 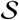. In this case we take the state to be *s* = (*x_s_, x*) and assume that the position of the searcher *x* is known to the agent, but the location of the source is not. As a consequence, the memory state is *m* = *b*(*x_s_*), where *b*(*x_s_*) is a probability map for the source location, and the memory space is 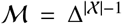 where 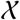 is the set of lattice points and Δ^X^ ‘ is the standard simplex. Actions are taken according to the current memory state and the memory is then updated following the Bayes theorem, making use of the known probability of observations *f(y**\**x, x_s_*) and transition probabilities *p*(*x*’|*x, a*), as pictorially shown in Fig. 4.

**Figure 4:**
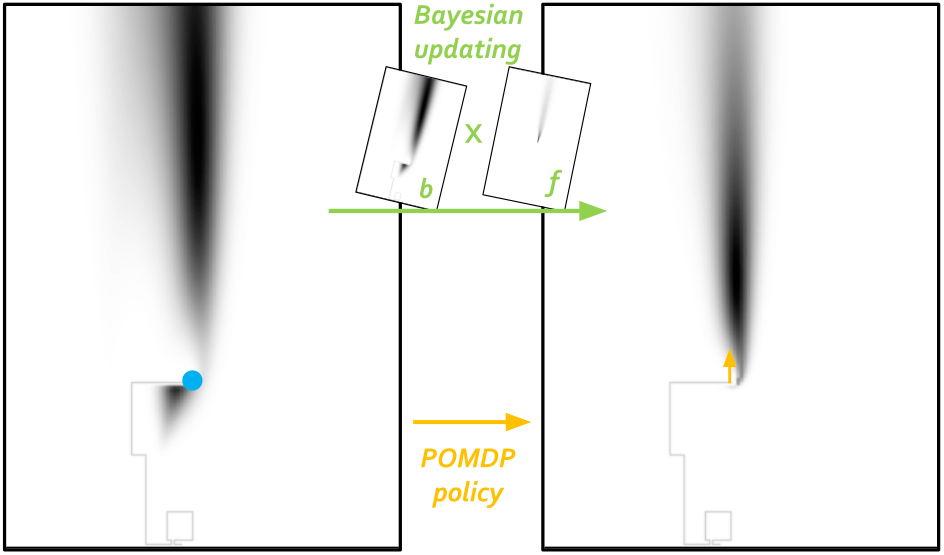
Graphical scheme of the Bayesian approach to POMDP. On the left, the memory of the past trajectory is encoded into the belief, shown as a grayscale heat-map. The instantaneous detection - shown as a blue filled circle - is used to update the belief according to the Bayes rule *b*(*x_s_*|*y*) ∝ *b*(*x_s_*)*f* (*y*|*x, x_s_*) and at the same time an action - in orange - is selected, depending on the current position *x* and the belief *b*(*x_s_*).

Bayesian POMDPs are inherently *model-based* as they use a priori knowledge of observation probabilities and transition probabilities in order to infer the state of the system. Leveraging on this knowledge, it is possible to perform a systematic optimization approach over the choices of actions, by approximating the solution of the Bellman’s equation with various methods [16, 17, 18, 19]. The computation of optimal actions is usually a daunting task so that one often makes recourse to heuristic approaches that often turn out to be very effective [13, 16, 20].

Furthermore, it is important to notice that in this setting the optimization is performed only over the choice of actions while the memory evolution is constrained to follow the Bayesian updating rules sketched in Fig 4 – it is, in this sense, hardwired. In the Bayesian approach the memory states are typically very high dimensional vectors, a fact that limits many applications. Moreover, if one wishes to consider continuous state spaces, the memory space becomes an even more unwieldy function space. A possible approach that reduces this complexity is to approximate the beliefs with Gaussian mixture models and combine it with some heuristic policy as in Ref.[21].

In the next subsection we will see an algorithm that attempts at a systematic optimization over memory updates as well and that can in general tackle continuous spaces as well.

#### 1.2.3. Recurrent neural networks: abstract memory spaces

The main motivation to use beliefs as memory states in Bayesian POMDPs is that they are a sufficient statistics for the full history of previous observations and actions. In this specific sense they are an optimal choice because they guarantee that no information about the past experience is lost. The price to pay is that the memory space is in general very large. It is natural to ask if more compact, abstract memory spaces can encode the past history without hugely sacrificing performance. The fact that the agent has a specific goal is likely of help in this situation, as one can hope that the information relevant for the task can actually be compressed in a lower-dimensional space. How can actions and new memory states be generated in this abstract setting? In the setting of model-based stochastic control, a possible approach has been outlined in Ref. [22], while in the model-free case an option is to parameterize the action generation and memory update process by means of a recurrent artificial neural network [23, 24] as depicted in Fig. 5. In the case at hand, the memory will be a vector in a space of suitable dimension, the inputs would be instantaneous detections *y_t_*, positions *x_t_* (and possibly the past action *a*_*t*-1_). As is apparent from the diagram, the neural network outputs at the same time actions and updated memory states. At any given time, the memory state of the network implicitly encodes the history of all past observations. The weights 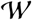 of the connections are free parameters that can be optimized in order to achieve better performances in terms of cumulative rewards. Thus, both memory updates and policy are optimized (or learned) simultaneously.

**Figure 5:**
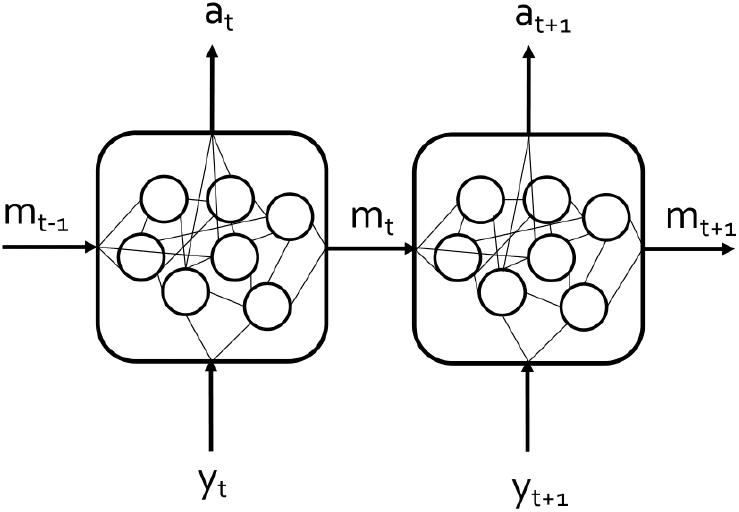
Time-unfolded recurrent neural network for POMDP. At each time t the past memory *m*_*t*_1_ and current observation *y_t_* are used to select an action *a_t_* and update the memory to *m_t_*. The weights are pictorially shown as the edges of the graph.

The advantage of using neural networks is that they can be straightforwardly used in much more complex environments than a discrete grid-world. They can be used with continuous states, actions, and observations, and are model-free: they are optimized, usually via Reinforcement Learning [15], by using data from experience (real or simulated) without relying on a priori knowledge of the environment. They can also gracefully deal with correlated observational inputs. Such an approach has been recently applied to olfactory search in a continuous environment [25].

There are also some disadvantages, though. First, optimization can demand large amounts of data. Second, the black-box character of the artificial neural network and the still sizeable vector space make the interpretation of the optimized behavior rather difficult. Typically, a substantial effort in post-processing and a further strong dimensionality reduction are required in order to make sense of the decisions that are made by the agent.

In the next section we introduce our approach in terms of finite-state controllers (FSC)[26, 27, 28, 29, 30, 31], which attempts at combining the positive aspects of the previous approaches while keeping the algorithm computationally inexpensive and easy to interpret (see Table 2). Finite-state controllers have discrete memory states which also provide an encoding of the history of past observations. However, at variance with recurrent neural networks, their interpretation is much more transparent, as we will show below.

**Table 2:** Comparison between algorithmic approaches.

| Algorithm | Memory | Model-based | Model-free | Policy | Memory update |
| --- | --- | --- | --- | --- | --- |
| Cast-and-surge | discrete | no | yes | hardwired | hardwired |
| Bayesian search | continuous | yes | no | optimized | hardwired |
| Recurrent neural network | continuous | no | yes | optimized | optimized |
| Finite-state controller | discrete | yes | yes | optimized | optimized |

## 2. Results

### 2.1. Finite-state controllers

As the name suggests, finite-state controllers use a memory that is a discrete set of finite cardinality, usually small. The controller dictates at the same time how to perform actions and update memories. One can think of these operations as a “super-action” that affects a “super-state” made by the pair (*s, m*) where the states *s* are only partially observable while the memories *m* are accessible to the agent [28]. In practice, to each memory-observation pair (*m, y*), the FSC associates a probability distribution *π_θ_* over actions and new memory states (*a, m*’). In other words, the controller is a map 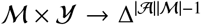. The probability distribution *π_θ_*(*a, m*’|*m, y*) completely defines the strategy of the agent, where *θ* is a set of parameters that can be optimized to obtain the best performance. For a POMDP finite-state controller, the maximum dimension of the parameter space is 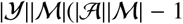). This value is attained if no specific graph structure is imposed a priori, i.e. the controller in principle allows all possible choices of memory updates and actions. Even in this case, this number is usually much smaller than the number of weights to be optimized for a recurrent neural network.

A graphical representation of a simple finite-state controller is given in Fig. 6.

**Figure 6:**
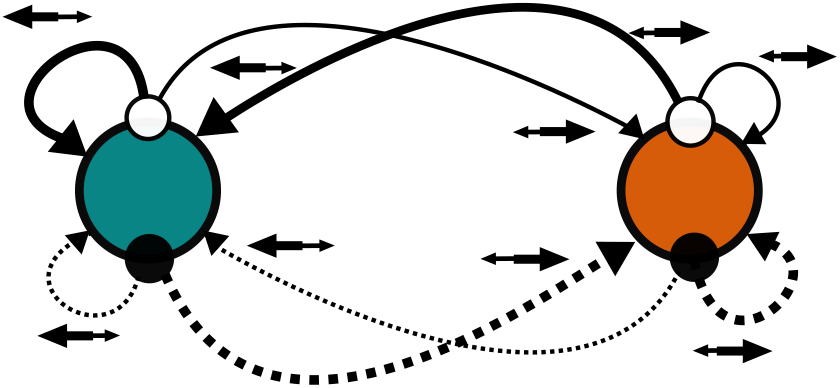
A graphical representation of a finite-state controller for a POMDP with two memories 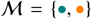, two observations 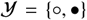 and two actions 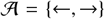. Memory updates are represented by continuous or dashed lines depending on the instantaneous observation. The thickness of line represent the probability of the corresponding transition. Associated actions (and their probability) are shown by the arrows (and relative sizes) above that transition. The transition probabilities and the probability of the associated actions are all parameters that are subject to optimization.

If the modality of the interaction between sensorimotor interface and agent is known – that is, if the agent knows the probability distribution of observations and new environmental states given the current environmental state – then it is possible to write closed and exact expressions for the gradient of the objective function (the expected cumulative reward) with respect to the policy parameters *θ*. The gradient can be efficiently computed if the search space is not exceedingly large (see Methods Sec 4) and optimization can be performed by straightforward gradient ascent. This approach is known as a *policy gradient* method [15]. In this sense, we can view finite-state controllers as belonging to the class of model-based, or planning, approaches such as Bayesian POMDPs. Under some conditions, policy gradient methods can attain global maxima whereas in general they are only guaranteed to reach local maxima [32].

What can be done when the agent does not know how the environment interacts with itself through the sensorimotor interface? Or when the state-space is so large that direct modelbased optimization is computationally unfeasible? In these cases it is still possible to *learn* the optimal POMDP controller by means of a data-driven approach, specifically a *stochastic policy gradient* method [15]. This method is model-free and it is indeed the one adopted to optimize recurrent neural networks – except that for finite-state controllers it is much less expensive on the computational level. While we discuss the model-free approach in the Methods Section, in the rest of the paper we will present results obtained by model-based, direct gradient search.

### 2.2. Optimizing finite-state controllers for olfactory search

We have created a virtual odor landscape using data from an experimental smoke plume used to study search behaviors in walking flies [6]. The probability of a detection is taken as the average fraction of time that the concentration signal at a point was above a chosen threshold. In order to check the robustness of the results we considered two cases, loosely corresponding to “weak” and “strong” odor regimes, with intensity at the source approximately 3× and 9× the threshold value, respectively. As for the search domain we have considered two different regions around the plume with different spatial resolution, corresponding to grids of size 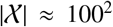. The results presented below are for the “strong” odor regime and the higher resolution case, but the main findings are similar for all cases (see Supplementary Information). The step-size has been chosen to be comparable to the typical scales observed in experiments with flies [6]. Searchers were initialized at the farthest downwind side of the domain and distributed along the crosswind distance with a probability proportional to the detection occurrence. Note that the agent receives information about the state *s* = (*x_s_, x*) only through odor encounters *y* ∈ {•, ⸰).

The memory 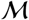 is a discrete set of cardinality 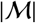 and we will focus on memory spaces of small size. The controller is parametrized by a softmax *π_θ_*(*a, m*’|*m, y*) ∝ exp(*θ_am’my_*). All transitions and action choices are possible a priori. Optimization is performed by Natural Gradient ascent (see Methods). Several, randomly chosen, initial choices of the parameters are used to mitigate the effects of trapping in local maxima.

We now turn to give a qualitative description of the behavior of optimized controllers (detailed quantitative results for strong and weak signals and for different spatial resolutions are given in the Supplementary Information). A comparison of the performances is given below.

#### Single memory: best reactive strategy

When there is a single memory state 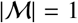 the agent can only make decisions based on the current observation, i.e. it follows a reactive strategy. In this case the optimal controller takes the form shown in Figure 7a. Hereafter for visualization purposes we identified the memory states with colors. Whenever a detection is made, denoted by the symbol •, a single upwind move is performed while in absence of odor (⸰) the agents perform an asymmetric random walk with a preference for up-down motion and a slight downward bias. This rudimentary search strategy is not very effective and in a large fraction of cases the agent is not able to find the source in the allotted time, see Table 3.

**Figure 7:**
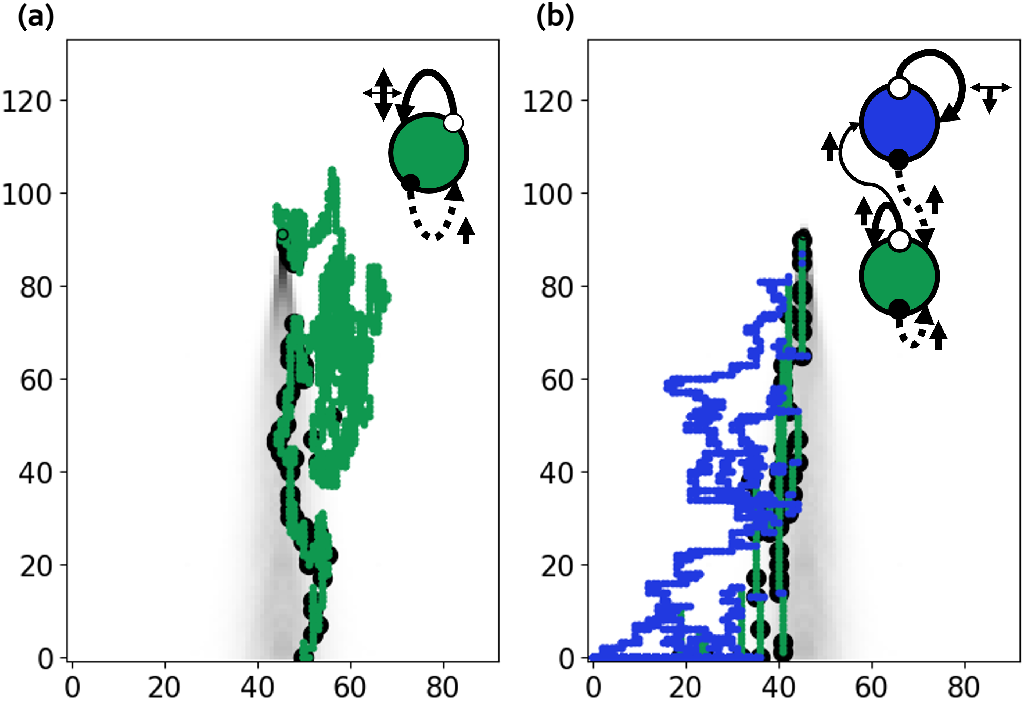
a) Graph of the optimal reactive controller 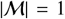 (top right) and a realization of a typical trajectory. b) Same for the optimal two-memory FSC 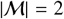. There are two observations 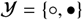 and four actions 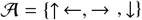. In this figure and the following ones, the gray shaded area is a map of the intermittency of the signal, defined as the fraction of time in which it is above the detection threshold (see Supplementary Information).

**Table 3:** Success rate and average search time for the algorithms mentioned in Table 2. Data obtained from 10^4^ samples. The performance indicators are computed for both cases of weak and strong signals. Average times are shown alongside their standard error. For reference, the minimum time to reach the source (if it were known to the agent), averaged over initial starting points, is 95.30.

|  | Algorithm | Success rate* | Average time** |
| --- | --- | --- | --- |
| Weak | Cast-and-surge | 81.53% | 460.6 ± 0.7 |
| | Infotaxis | 99.80% | 318.0 ± 1.8 |
| | 1M-FSC | 57.11% | 3047.6 ± 32.5 |
| | 2M-FSC | 95.69% | 1903.5 ± 23.2 |
| | 3M-FSC | 99.94% | 1602.5 ± 14.7 |
| | 4M-FSC | 100% | 983.0 ± 9.3 |
| Strong | Cast-and-surge | 64.08% | 309.6 ± 0.5 |
| | Infotaxis | 99.65% | 206.5 ± 0.6 |
| | 1M-FSC | 68.39% | 2450.2 ± 28.7 |
| | 2M-FSC | 97.7% | 1603.5 ± 21.1 |
| | 3M-FSC | 100% | 942.1 ± 9.2 |
| | 4M-FSC | 100% | 690.6 ± 6.4 |

\* The search is successful if the source is found in less than $10^4$ steps.
\*\* Average time is computed only over successful searches.

#### Two memory states: surge and backward random walk

The presence of a second memory state (see Fig. 7b) leads to a more structured strategy. First, in absence of odor detections (⸰) surges can now persist for a geometrically distributed time, with a mean time 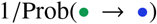. Second, the movement pattern associated with the blue memory 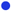) is a backward biased random walk with a larger probability of left and right moves. Third, upon odor detection (•), the memory is always reset to the green state, and a one-step upwind move is performed (dashed line transitions). The combined effect is a more effective plume recovery and faster upwind search, leading to a better success rate (see Table 3) and shorter average search time with respect to the reactive strategy.

#### Three memory states: emergence of a surge-and-cast pattern

For three memory states, the optimal controller takes the form shown in Figure 8 (see also the video of a trajectory in a dynamic plume). As for the two-memory case, upon odor detection, the memory is always reset to the green state, and a one-step upwind move is performed (dashed lines). In absence of odor (⸰) memory updates are in general stochastic (full line transitions). If the memory is in the green state, the most likely event is to stay in that memory state, and with a relatively small probability to transition to the yellow memory. As a result, the residence time in the green state is geometrically distributed with average equal to 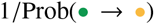. In plain words, after a detection, the agent performs an upwind surge with an average duration that has been determined by the optimization. Once in the yellow memory state, the agent stays there for a certain time, performing a cast to the right (looking upwind), with some random backward component of motion. The persistence in the yellow state is given by 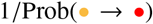. If detections are still absent, eventually the system transitions to the red memory and performs a cast to the left. In case of a prolonged period without odor encounters, the agent alternately casts left and right and slowly recedes downwind, a behavior that is instrumental in recovering contact with the odor plume.

**Figure 8:**
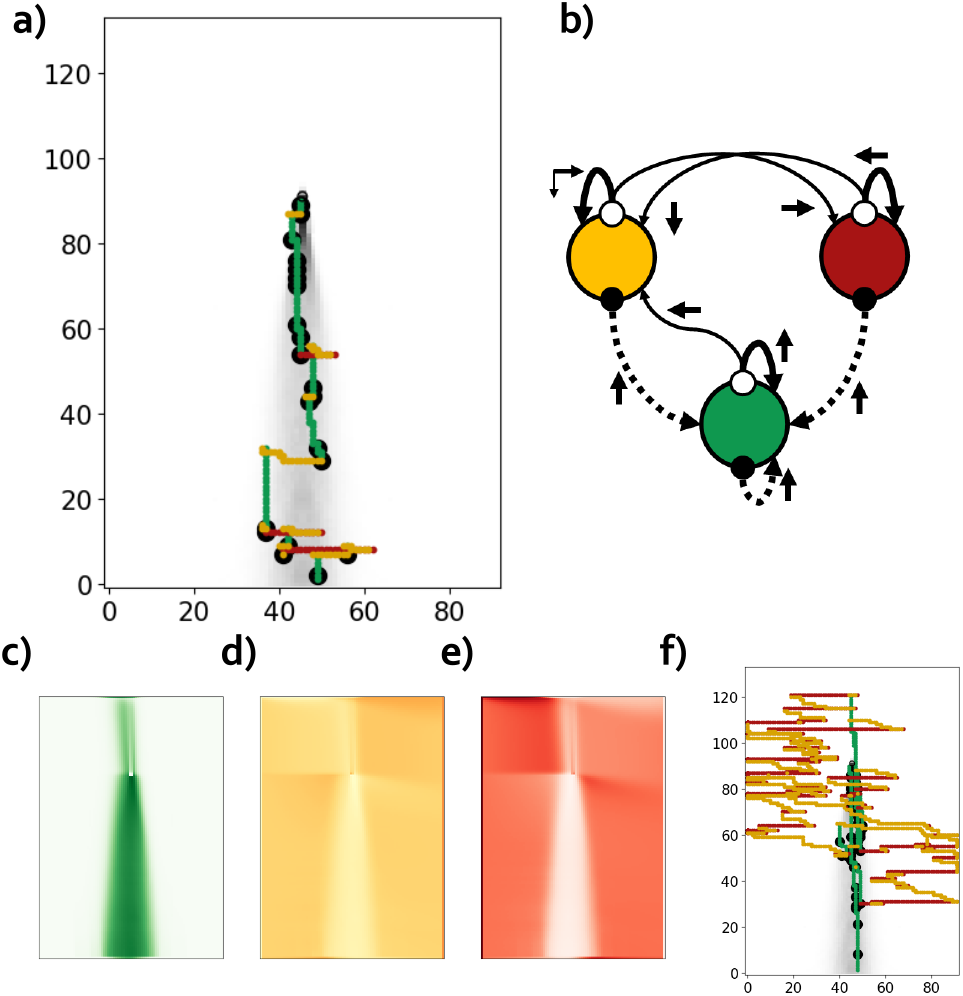
The optimal controller with 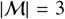 memories discovers a stylized surge/cast strategy. a) A typical trajectory, colored according to the memory state, detections are shown as •. b) Graph of the FSC, with the notation de-scribed in Fig.6. c,d,e) Heat maps for the spatial memory occupation Prob(*m*|*x*). Notice that the presence of boundaries leads to an increased probability of being in the red memory on the left side of wind axis, and vice versa. f) Example of a long trajectory. The initial overshoot (green portion of the trajectory upwind of the source) is compensated by repeated casts with a downward drift. All panels follow the same color coding for the memories.

Further insight in the interpretation of the abstract memory states can be gained by looking at Prob(*m*|*x*), the probability of being in a memory state *m* when the searcher is in *x*. As shown in Figure 8, the occupancy of the green memory is mostly localized downstream of the source, whereas the yellow and red memory are largely visited when the agent is away from the wind axis. Altogether, memory states can then be understood as coarse-grained beliefs according to the dictionary: 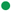 = within the plume, 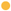 or 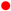 = outside the plume.

The process generated by the FSC is evidently asymmetric in its choice of actions, even if, by construction, the detection and transition probability do not break the left-right symmetry about the wind axis. Symmetry is spontaneously broken by optimization: performing the gradient ascent with a symmetric initialization inevitably leads to asymmetric solutions such as the one displayed here, just because of a small noise in the gradient evaluation. Incidentally, we observed that the reset memory state always coincides with the memory chosen to initialize the search (whose choice breaks the permutation symmetry under relabeling of the memories).

#### A locally optimal, globally suboptimal, three-memory controller: surge, up-down search, and backward-biased random walk

Gradient ascent also finds local optima, yet globally sub-optimal, whose controllers have an interesting behavior. In Figure 9 we show a controller that behaves as an approximate clock, where surges are followed by backward-biased random walks. Similarly as for the optimal controller, the green state 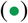 is associated with surges, and when detections are made (•), the controller is reset to that state (dashed arrows). In absence of detections (⸰), the surging behavior is kept for a geometrically distributed time and then a transition to the pink state 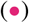 occurs. In this state, the agent performs a downward-biased random walk along the mean wind direction. If the agent still does not make contact with the odor, then the controller transitions to the purple state 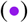 where it performs another kind of random walk but now with a dominant crosswind component and some backward bias, leading to a downwind zigzagging pattern. The performance of this FSC is ≈ 30% less than the optimal one.

**Figure 9:**
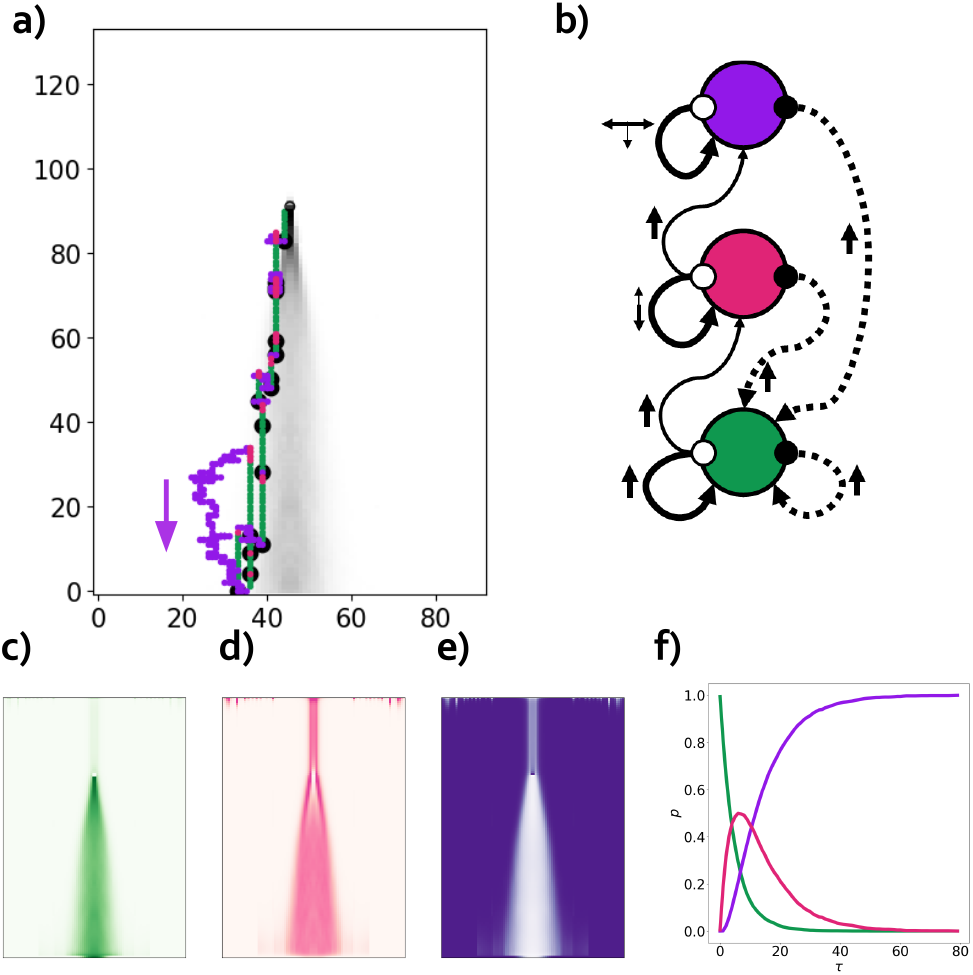
Second-best controller with 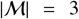 memories, displaying a surge/up-down search/backward random walk strategy. a) A typical trajectory. b) Graph of the FSC. c,d,e) Heat maps for the spatial memory occupation Prob(*m|x*). f) Occupation of memories Prob(*m|τ*) as a function of *τ*, the time since last detection. All panels follow the same color coding for the memories.

**Figure 10:**
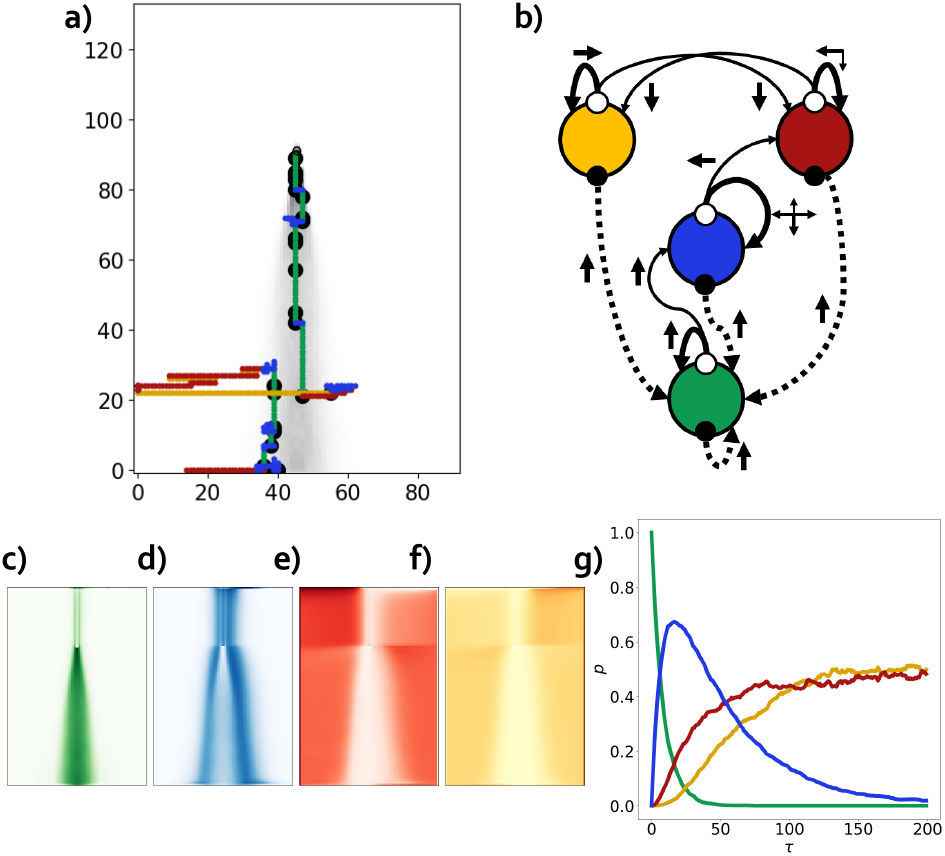
The optimal controller with 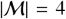 memories. a) Typical trajectory. b) Graph of the FSC. c,d,e,f) Heat maps for the spatial memory occupation Prob(*m|x*) g) Occupation of memories Prob(m**|**τ) as a function of time since last detection τ. All panels follow the same color convention for the memories.

It is natural to interpret these behaviors as successive efforts in reestablishing contact with the odor plume. This is also visible by measuring the probability Prob(*m|τ*) of being in a memory state conditioned to the time elapsed since the last detection τ, see Fig. 9f. The search along the wind axis in the pink state can be seen as a first recovery strategy while the purple state implements a last ditch attempt. The inspection of Prob(*m|x*) reveals that these are activated in different portions of space (Fig. 9 panels c, d and e). The green state associated with surges is essentially active inside the plume. The pink up/downwind search is active at the edge of the plume and straight upwind of the source, where it clearly represents an effective recovery search strategy. The purple state, which produces the downwind zigzagging, is the most active outside of the regions of activity of the other two states, where motions occurring only along the wind axis are quite ineffective.

#### Four memory states: richer behavioral patterns combine modules from smaller memory sets

The optimal controller with four memory states appears to compound features of both the optimal and sub-optimal three-memory controllers (see Fig. 8 and the video of a trajectory in a dynamic plume). Out of the four memory states, three have an analogous function as in the optimal 3M-FSC and are therefore similarly colored as green, yellow and red. These three memories are associated with surging 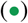 and left/right casting 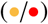. When the controller is in the fourth memory state 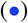 the agent performs a biased random walk that combines the behavior of the pink and the purple memory states in the suboptimal 3M-FSC of Fig. 9. Notice that while transitions between red and yellow are allowed in both directions, there is no transition from neither yellow nor red back to the blue memory. As seen for previous controllers, the detection of a signal acts as a hard reset: the system always transitions to the green state and restarts with an upwind move.

As previously, memory states can also be interpreted as coarse maps of the search space, where 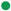 = within the plume, 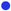 = on the edges of the plume or upwind of the source, and 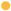 or 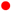 = outside the plume. Again, the transition diagrams shows properties akin to an approximate clock which in absence of detections follows the sequence 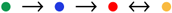, and is reset to the green state upon an odor encounter.

#### 2.2.1. Performance of optimal finite-state controllers

As shown in Table 3 the performance of the optimal finite-state controllers is remarkable given their simplicity. Hardwired cast-and-surge has large failure rates– likely because of the absence of an effective downwind recovery strategy. By the same criteria, even Infotaxis, a well-known heuristic Bayesian search strategy [20], falls short of a full success rate, if by a small amount, because of occasional pinning of trajectories around a location where an unexpected detection has occurred. FSCs with sufficient memory states achieve a 100% success rate, probably by virtue of their more exploratory character, albeit at the expense of an increase in search time. Indeed, the four-memory FSC searcher is about three times slower than Infotaxis to reach the source. Given that the memory space has been compressed from the belief space (a simplex of dimension 10^4^) to a discrete set with just four elements, this loss in performance seems excusable. As a side remark, we do not show simulations with recurrent neural networks because performances heavily depend on the choice of the architecture of the network and it is therefore very difficult to design a meaningful term of comparison with other algorithms.

### 2.3. Temporal abstraction in optimal finite-state controllers

A shared feature of the controllers described above is the resetting to the state 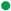 whenever a detection (•) is made, and the same action (↓) is chosen irrespective of the memory. As a result, after an odour encounter and until the next one – that is during each “blank” according to the terminology of Ref. [7] – the searcher enters a stochastic program entirely defined by the FSC graph. Any information other than the time since the last detection is not required for its execution. This program has a hierarchical structure with two separate time-scales: a slow one, over which transitions between memory states *m* → *m*’ occur, and a fast one associated with actions a. Each memory state can then be interpreted as a high-level “option” (see Ref. [33]) that induces a specific time-extended course of low-level primitive actions (e.g. “surge”, or “cast”, etc). The finite-state controller orchestrates the sequence of options, until, eventually, a new odor encounter occurs and the program starts entirely afresh.

## 3. Conclusions and perspectives

In this paper we have shown how simple finite-state controllers can be optimized to discover effective strategies for olfactory search. The resulting patterns of motion of the searcher are easy to interpret and bear a strong resemblance to behaviors that have been actually observed in living organisms. Besides their biological relevance, algorithms for olfactory search are also of obvious interest for robotics [34, 35]: in this respect FSCs may offer an alternative with very low computational load.

We focused on small memory spaces with the aim of identifying minimal models, at the expense of performance. Is this approach scalable [29]? Doubling the size of the FSC requires in general a fourfold increase in the space of parameters to be optimized, which quickly leads to difficulties. However, if a specific structure of the memory transitions is imposed – e.g. a tree-like graph – then FSC with larger memory sizes could be approachable. Also, based on our observation that optimal controllers with larger memories are actually compositions of smaller ones, one can hope that effective FSC could be built by assembling simpler modular structures.

We have offered a proof of principle of the usefulness and versatility of FSC algorithms in an admittedly simple virtual odor environment. In order to apply the method to more realistic settings, one needs to fully embrace model-free optimization. As explained above (see also Methods), FSC can be optimized by stochastic gradient methods that do not require extensive model knowledge and lend themselves to be used in continuous, complex search spaces. However, these algorithms are often plagued by high variance and require a careful use of baselines and other algorithmic devices: more work is needed in this direction as well.

It remains an open question whether these abstract memory states have any significance for the neuroscience of olfactory search (see e.g. [36] and the discussion in Ref. [25]). The emergence of a hierarchical structure in the optimal finite-state controllers could be of help in identifying neural correlates [37] while the resetting property hints at a relationship with time-coding neurons [38]. We expect that a close comparison between finite-state controllers and neural models could shed further light on the algorithmic underpinnings of search and navigation behaviors in living organisms.

## 4. Methods

### 4.1. Policy-Gradient method for FSC

We show here how to optimize the parameters of the finite-state controller using the model-based policy gradient.

We will use standard results for policy gradient [15], applying them to state-memory pairs (*s, m*) and action-memory update pairs (*a, m*’). Recall that *s* = (*x, x_s_*) where *x* is the agent’s position and *xs* is the source location. Both are unknown to the agent. We assume memory transition to be flawless, i.e. when a memory transition is selected, it occurs with probability one.

An exact expression for the policy gradient can be obtained by using the known transitions between different combined states *T*(*s*’, *m*’ *s, m*). This is true even if the agent does not have access to the information about the spatial terms *s*’ and *s*. The resulting gradient for the policy will be, however, a weighted contributions from all state-memories pair *s, m*, see Eq.7. In detail, the transition between state pairs can be written as

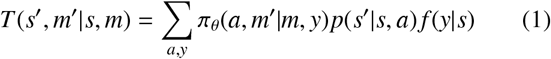

where *π_θ_* is the parameterized policy which selects spatial action *a* and memory update *m*’, and all the other quantities are defined as in Table 2. The reward function is equal to -(1 - *γ*), where 0 ≤ *γ* < 1 is the discount factor, except when *x*’ is the target location, in which case it is zero. The average reward *r*(*s, m*) is then defined as

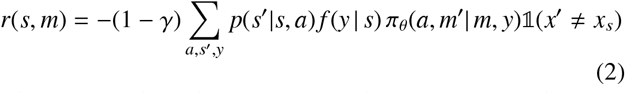

where 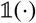 is the indicator function that equals one if the argument is true, and zero otherwise. Given that the initial position of the agent is randomly chosen following a distribution *ρ*, the solutions for the average occupancy of the state-memory pair reads, in matrix form

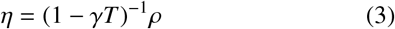

The value function *V*(*s, m*) of the policy is similarly given by

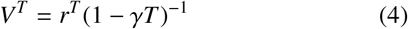

Where *A^T^* denotes the transpose of *A*.

The optimization process corresponds to maximizing the expected return

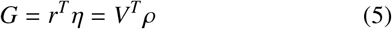

which by definition is comprised between −1 and 0. For *γ* → 1 one has *G* ≈ -(1 - *γ*)× (average time of search).

For the policy, we use a Boltzmann parameterization

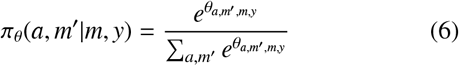

The (natural) gradient is given by the policy-gradient theorem [15, 39], which in this case reduces to

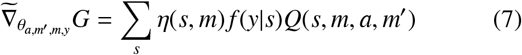

where *Q*(*s, m, a, m*’) is related to the value function introduced above by

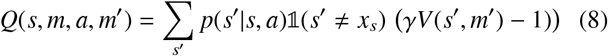

Notice that the gradient depends on the initial distribution *ρ* through the occupancy *η*. We see that in order to find the (natural) gradient 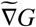, one needs to solve the two linear equations (3) and (4) which return the value function *V*(*s, m*) and the average occupancy of the state-memory pair *η*(*s, m*).

#### 4.1.1. Iterative solution

The solution of equations (3) and (4) requires the inversion of a large matrix. As the size of the domain 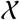 increases, this becomes the computational bottleneck of the gradient computation. To overcome this issue, we employ the Jacobi iteration method. The average occupancy η can be found as the asymptotic value of the iteration

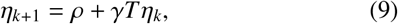

for *k* → ∞. Similarly, *V* is calculated via

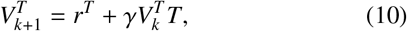

for *k* → ∞. The sequence is guaranteed to converge, since *γ* < 1 and the modulus of the largest eigenvalue of *T* is at most one thanks to the Perron-Frobenius theorem.

### 4.2. Model-free Policy Gradient

It is not strictly necessary to know *p*(*s*’ |*s, a*) and *f*(y|s) in order to estimate the gradient. And even if these objects are known, it might be too expensive or virtually impossible to perform the exact evaluation of the gradient if the state space is too large. Instead, one can use the stochastic, model-free version of the policy-gradient theorem to approximate the true (natural) gradient:

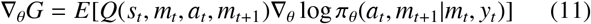

In this case one can generate stochastic trajectories of the search by the agent, record the sequences of tuples *m_t_, y_t_, a_t_, r_t_* and estimate 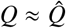 - e.g. via Monte Carlo methods. The estimator of the natural gradient then reads

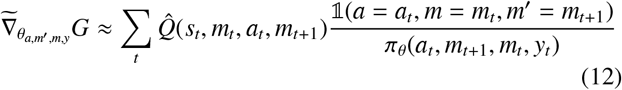

where the sum is performed along the trajectory. This estimator is unbiased but typically has a large variance, so that bootstrapping and variance reduction techniques are customarily used [15].

## Acknowledgments

This project has received funding from the European Union’s Horizon 2020 research and innovation programme under the Marie Skłodowska-Curie grant agreement N°956457.

