## Supplementary Information for "Olfactory search with finite-state controllers"

### Supplementary Information:
Olfactory search with finite-state controllers

Kyrell Vann Verano[a,b], Emanuele Panizon[a], Antonio Celani[a,*]

[a] *The Abdus Salam International Center for Theoretical Physics, ICTP, Strada Costiera 11, 34151 Trieste, Italy*
[b] *Dept. of Physics, Università degli Studi di Trieste, Via Alfonso Valerio 2, 34127 Trieste, Italy*

#### 1. A virtual odor environment from experimental plume data

The data used in this section are taken from Ref. [1, 2]. where a wind-tunnel assay for the simultaneous measurement of *Drosophila melanogaster* behavior and odor concentration was developed. The concentration of odor (in this case smoke, which attracts flies) was recorded into two-dimensional videos of intensity $c(x, t)$. We have processed the instantaneous concentration fields so as to obtain a stationary probability of detection as follows. We set the probability of a detection (•) at a position $x$ as the average fraction of time in which the signal was above an intensity threshold $w$ (also called "intermittency")

$$p(\bullet|x) = \frac{1}{T}\sum_{t=1}^{T} \mathbb{1}(c(x,t) > w). \tag{1}$$

We considered two different regions of interest at different resolution and two different concentration thresholds $w$. Since actions are taken at each step, and considering the size of the grid and the average speed of a fly, the time between each step was taken to be approximately consistent with the typical frequency of changes in behavior of the flies as reported in [1]. Figure 1 summarizes the data processing pipeline.

#### 2. Infotaxis

Here we briefly summarize the Infotaxis heuristic algorithm [3]. The episodes are initialized with the agent in a position $x \sim \rho(x)$, compatible with the initialization of FSC searchers, and a flat belief $b_0$ for the target position $x_s$. Note that in other works a different initialization, enforcing a detection at the beginning of the episode, is chosen [4].

At each time-step, the algorithm evaluates the expected entropy decrease in the belief following the choice of an action $a$. This is equal to the mutual information $I(X_s : Y|A = a, X = x)$ between the observation $Y$ and the location of the target $X_s$, given the position of the agent $x$.

The expected entropy decrease contains multiple terms. The first is related to the probability of finding the target in $x'(a)$, the state reached after taking the action $a$, which will reduce the entropy of $b$ to zero. The other terms correspond to the event of *not* finding the target and receiving an observation $y$, with a probability $p(y|x'(a), b) = \sum_{x_s \neq x'(a)} b(x_s) f(y|x'(a), x_s)$. As a consequence of the observation, the belief is updated via Bayes' rule:

$$b'_{|a,y}(x_s) = \frac{b(x_s) f(y|x'(a), x_s)}{\sum_{x_s} b(x_s) f(y|x'(a), x_s)} \tag{2}$$

Summing all contributions one has

$$I(X_s : Y|A = a, X = x) = -b(x'(a))H(b) + \sum_{y} p(y|x'(a), b)(H(b_{|a,y}) - H(b)) \tag{3}$$

*Corresponding author

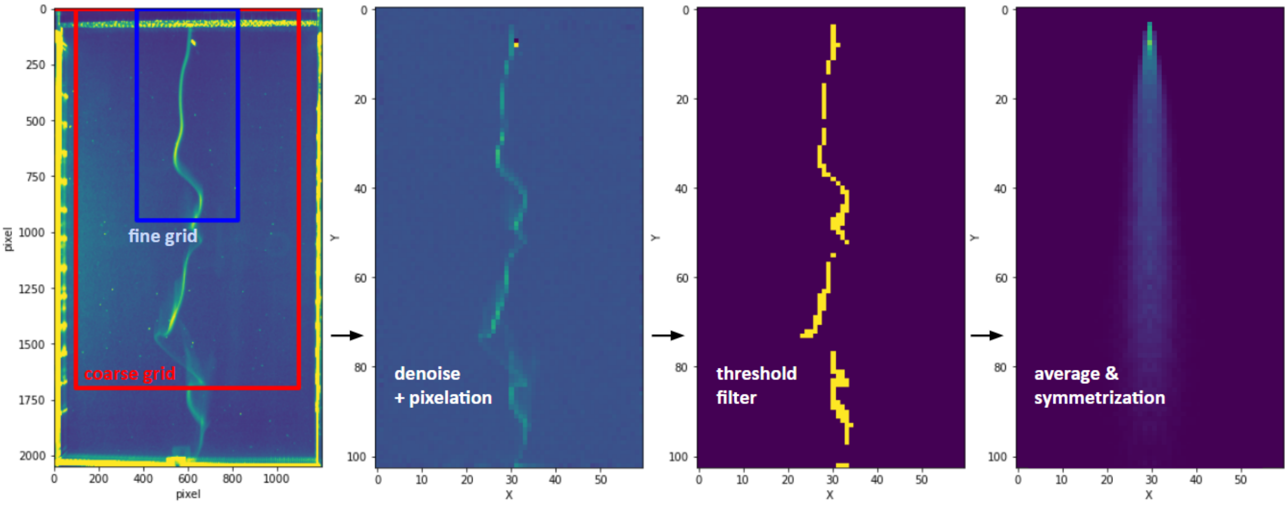


Figure 1: Processing of experimental plume data. From left to right. i) A frame of the original video. The range of the two different grids are shown by the blue and red rectangles. ii) The effect of removing background noise and pixelation. iii) Regions where the signal intensity is above a certain threshold. iv) Average over 8500 frames, after enforcing symmetry along the wind axis, to avoid spurious asymmetry effects on the policy of the finite-state controller.

The Infotaxis strategy then prescribes to take the action

$$a = \mathrm{argmax}_a I(X_s : Y|A = a, X = x) \tag{4}$$

until the target is found.

As discussed in Refs. [4, 5] the performance of infotaxis is quite close to the *bona fide* optimal Bayesian POMDP solution.

#### 3. Finite State Controllers

In this section we present the detailed results for Finite-State Controllers (FSC) with $|\mathcal{M}| = 1, \dots, 4$ memory states, both for weak and strong signals (two different thresholds) and the two spatial resolution levels (fine grid and coarse grid). In each figure we present (a) $\pi(a, m'|m, y = \circ)$, the probabilities of taking action $a$ and updating to memory $m'$ from $m$ whenever there is no detection (b) the graph of the FSC of memory updates and actions taken, (c) the distribution of search times, and (d) a sample trajectory. As mentioned above there is a hard reset to the memory $\bullet$, so $\pi(a = \uparrow, m' = \bullet|m, y = \bullet) = 1$. While the structure of the FSC with a given memory size $|\mathcal{M}|$ remains the same for weak/strong and fine/coarse cases, the transitions probability change, reflecting different optimal durations of behavioral patterns. For instance, in a weak signal regime or in coarse grid, the searcher tends to do longer surges. Another interesting difference is observed in the weak/coarse case (see Figure 11) where the backward random walk is replaced by another surging upwind instead. As shown in the trajectory (Figure 11d) the searcher finds the target even if only three detections are made.

In Figure 12 we show the results of optimization from different initial conditions, showing the existence of suboptimal *bona fide* extrema.

#### 4. Sample videos of FSC in the dynamic plume environment

Sample trajectory with 3 Memories: Video1 Sample trajectory with 4 Memories: Video2

*Acknowledgments*

This project has received funding from the European Union's Horizon 2020 research and innovation programme under the Marie Skłodowska-Curie grant agreement N°956457.

| | Algorithm | Success rate* | Average time** |
|---|---|---|---|
| Weak | 1M-FSC | 28.53% | 93.5 ± 0.1 |
| | 2M-FSC | 88.80% | 1348.7 ± 21.8 |
| | 3M-FSC | 98.56% | 1676.5 ± 18.2 |
| | 4M-FSC | 99.42% | 1327.4 ± 15.6 |
| Strong | 1M-FSC | 73.74% | 2160.2 ± 25.8 |
| | 2M-FSC | 97.86% | 1568.9 ± 20.7 |
| | 3M-FSC | 100% | 717.1 ± 7.5 |
| | 4M-FSC | 100% | 498.5 ± 5.0 |

Table 1: Success rate and average search time for the algorithms in the case of a *coarse grid*. In the main text we showed results for the fine grid. Data obtained from $10^4$ samples. The performance indicators are computed for both cases of weak and strong signals. Average times are shown alongside their standard error. For reference, the minimum time to reach the source (if it were known to the agent), averaged over initial starting points, is 93.50.

* The search is successful if the source is found in less than $10^4$ steps.

** Average time is computed only over successful searches.

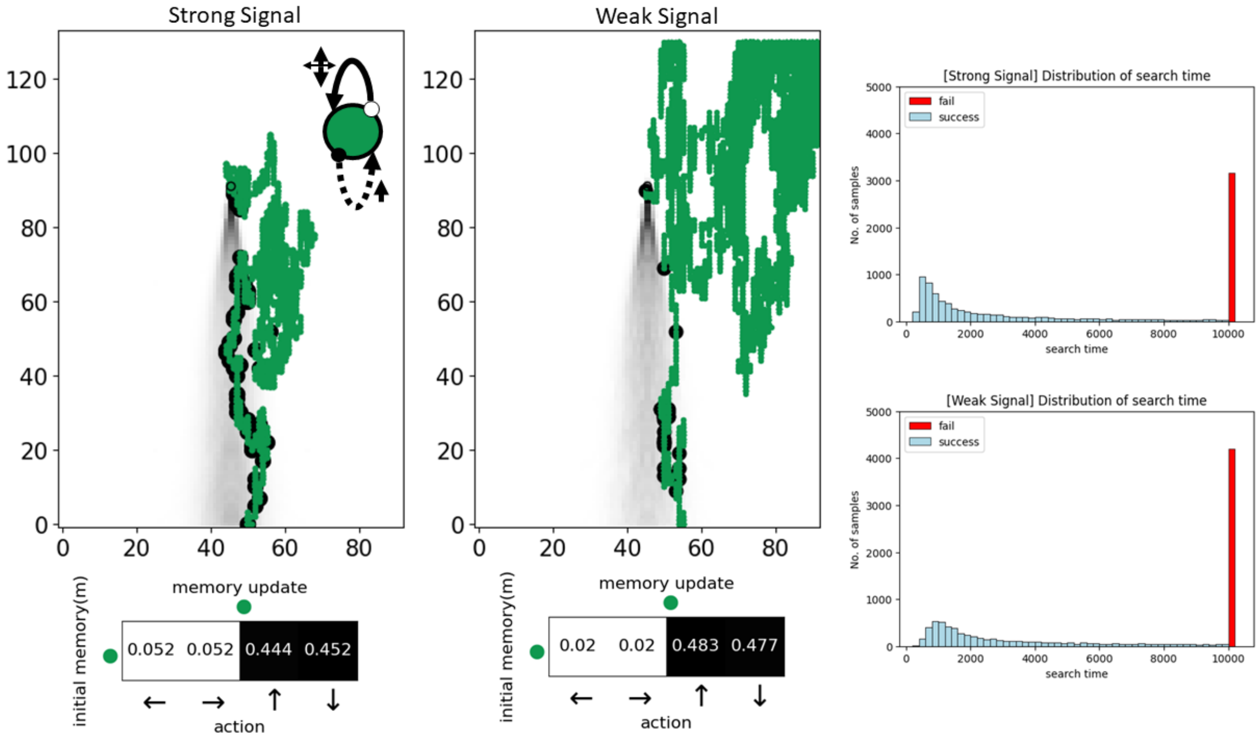


Figure 2: Finite state controllers with $|\mathcal{M}| = 1$ in a fine grid.

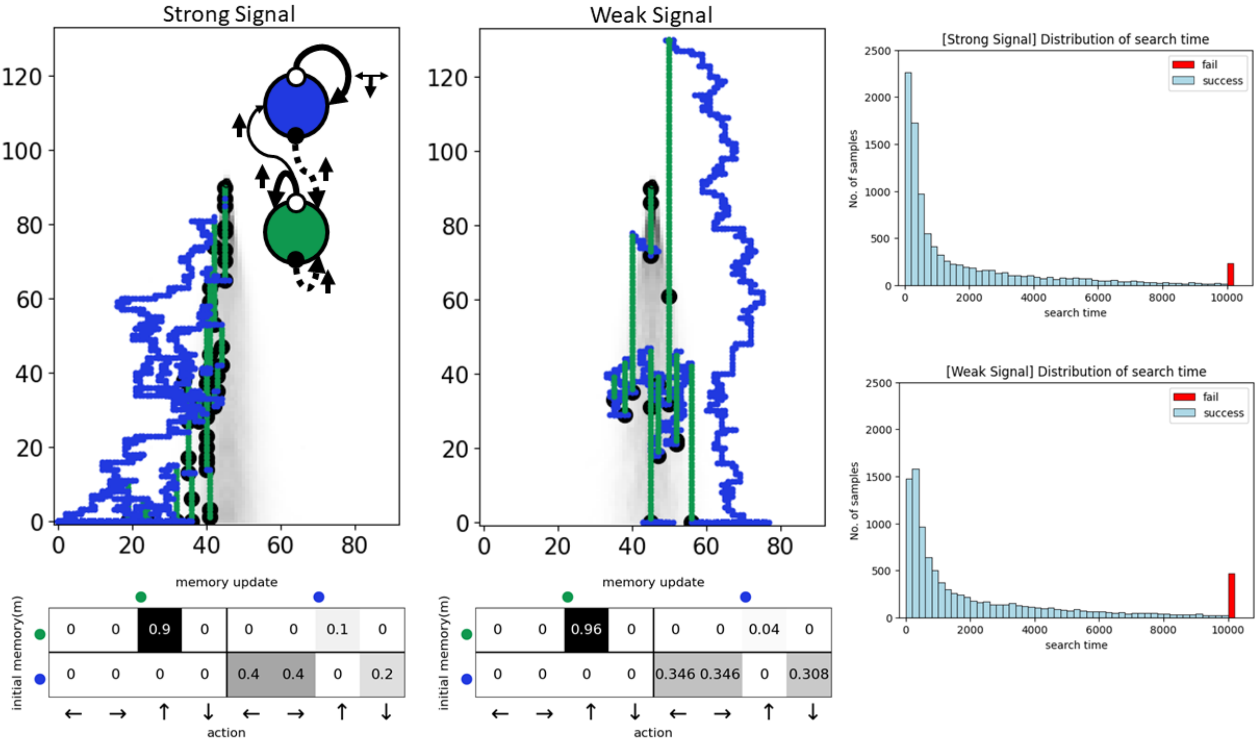


Figure 3: Finite state controllers with $|\mathcal{M}| = 2$ in a fine grid

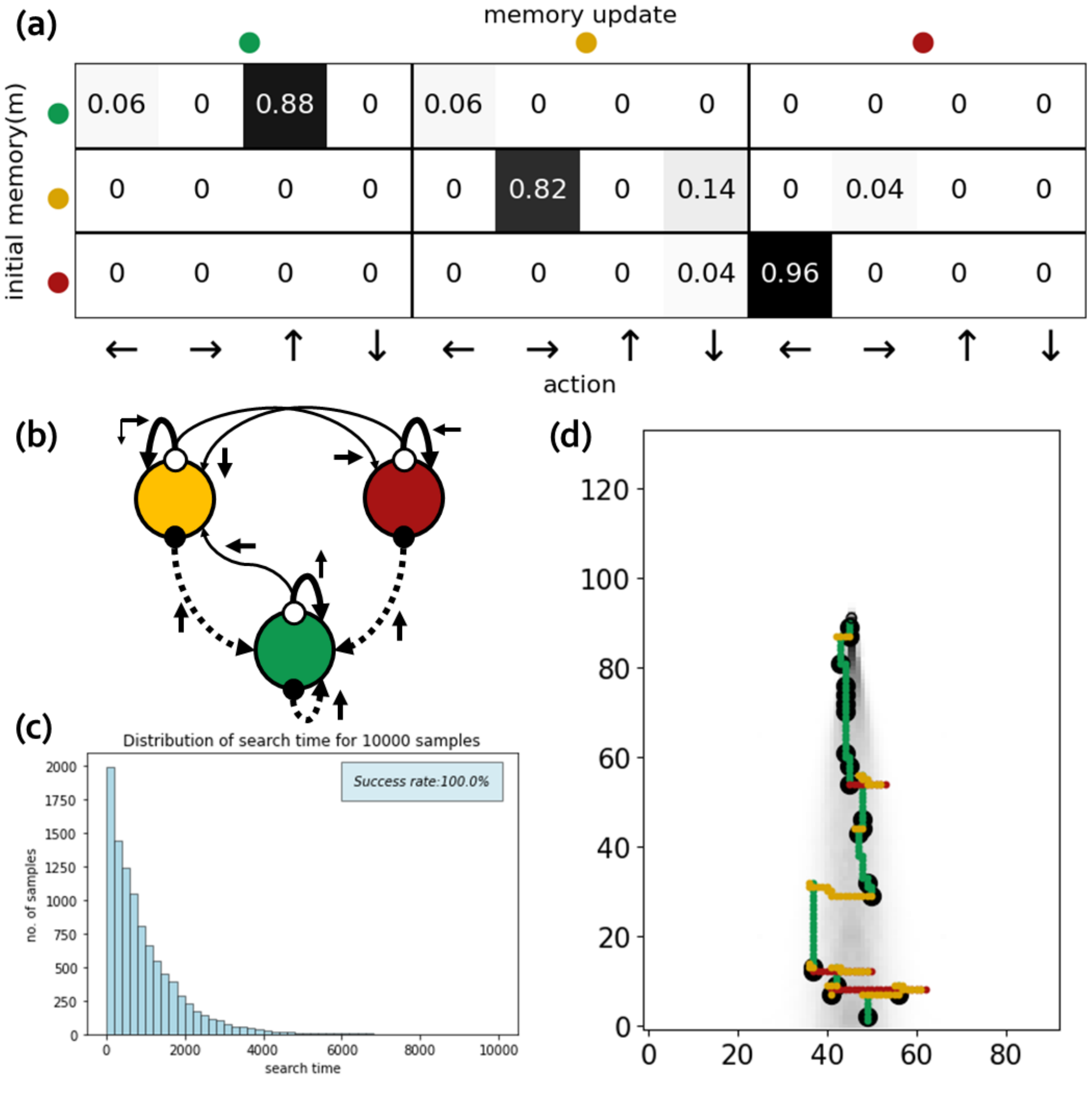


Figure 4: Finite state controllers with $|\mathcal{M}|$ =3 in a fine grid with strong odor signal

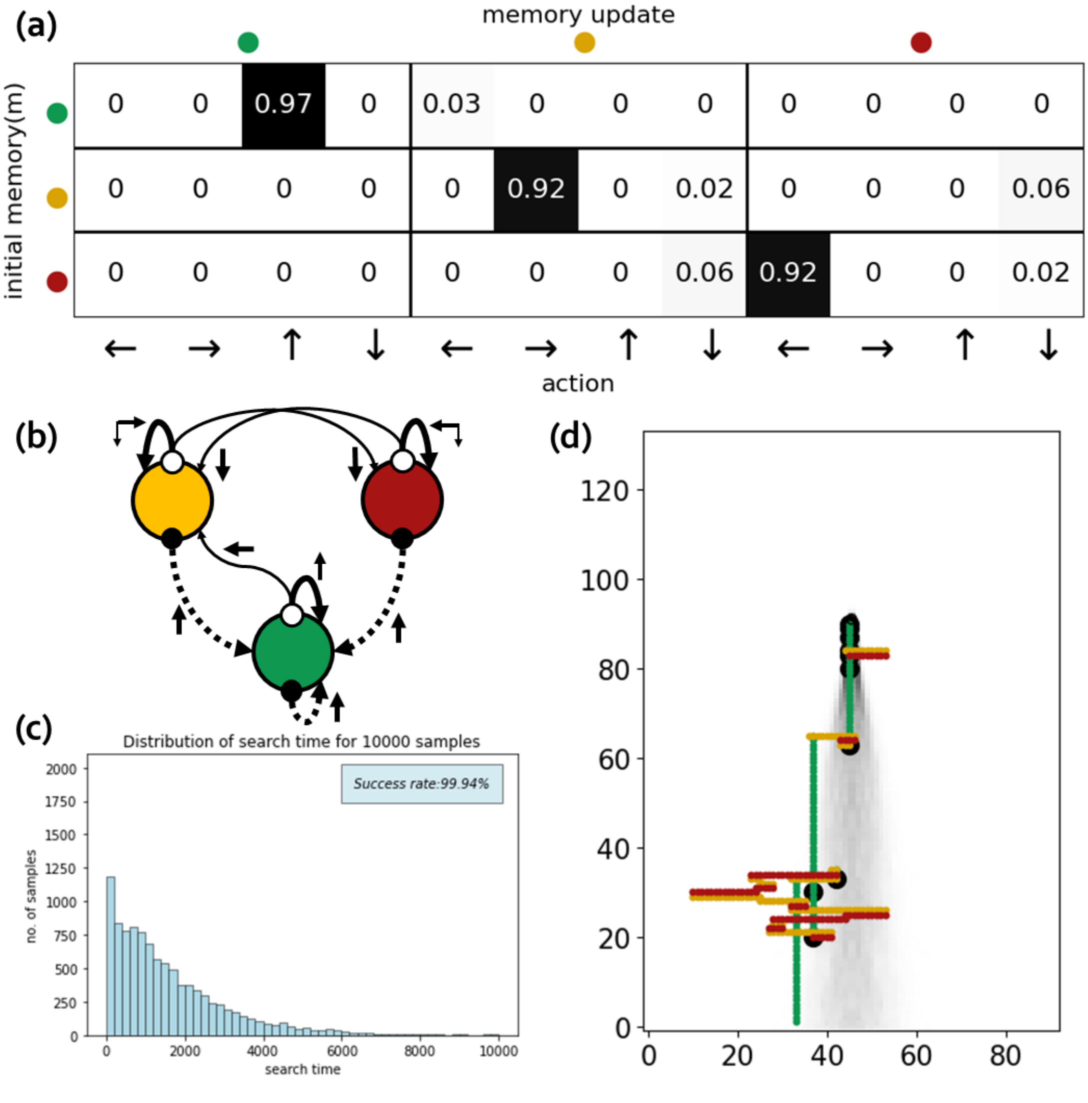


Figure 5: Finite state controllers with $|\mathcal{M}|$ =3 in a fine grid with weak odor signal

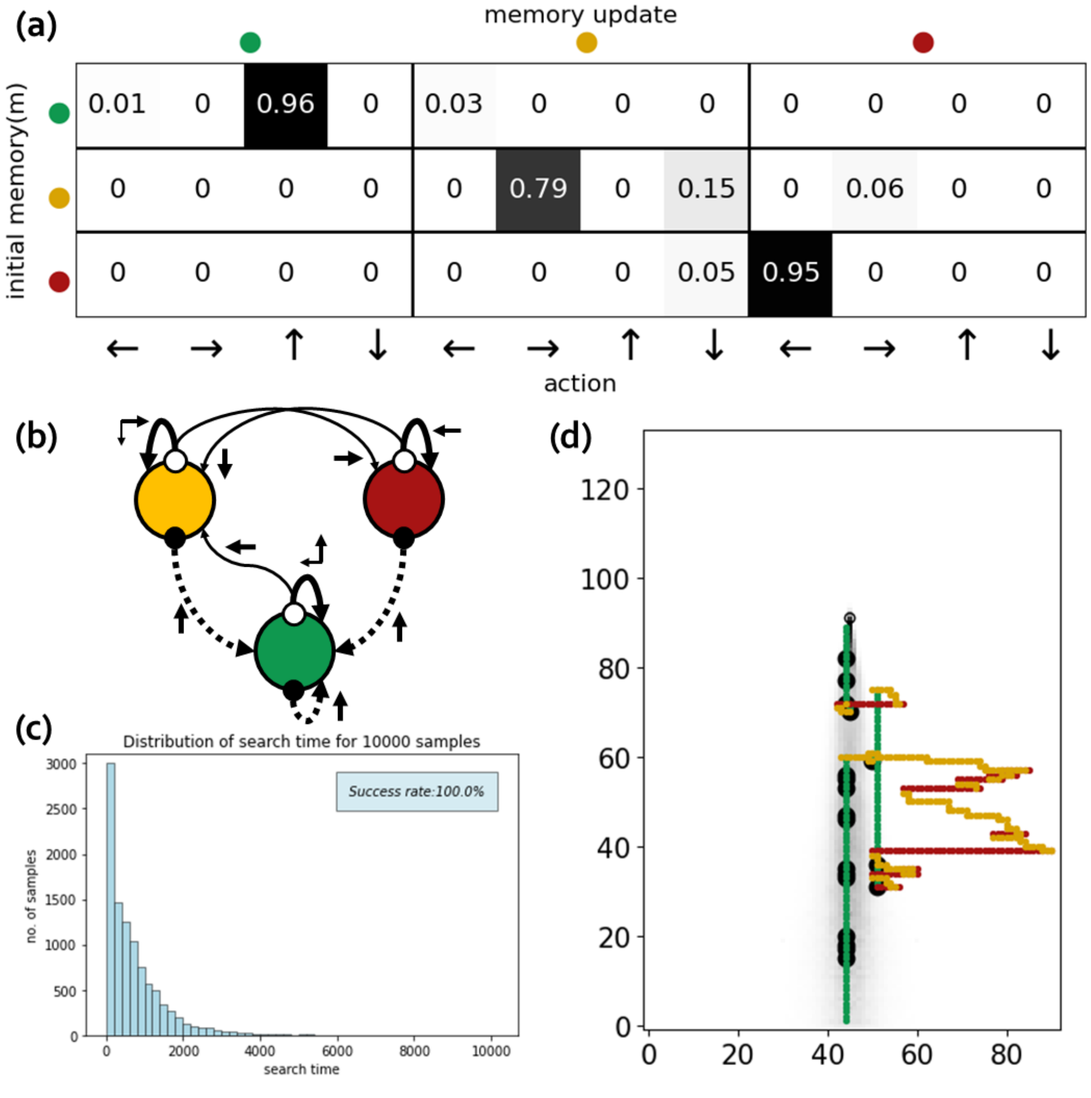


Figure 6: Finite state controllers with $|\mathcal{M}|$ =3 in a coarse grid with strong odor signal

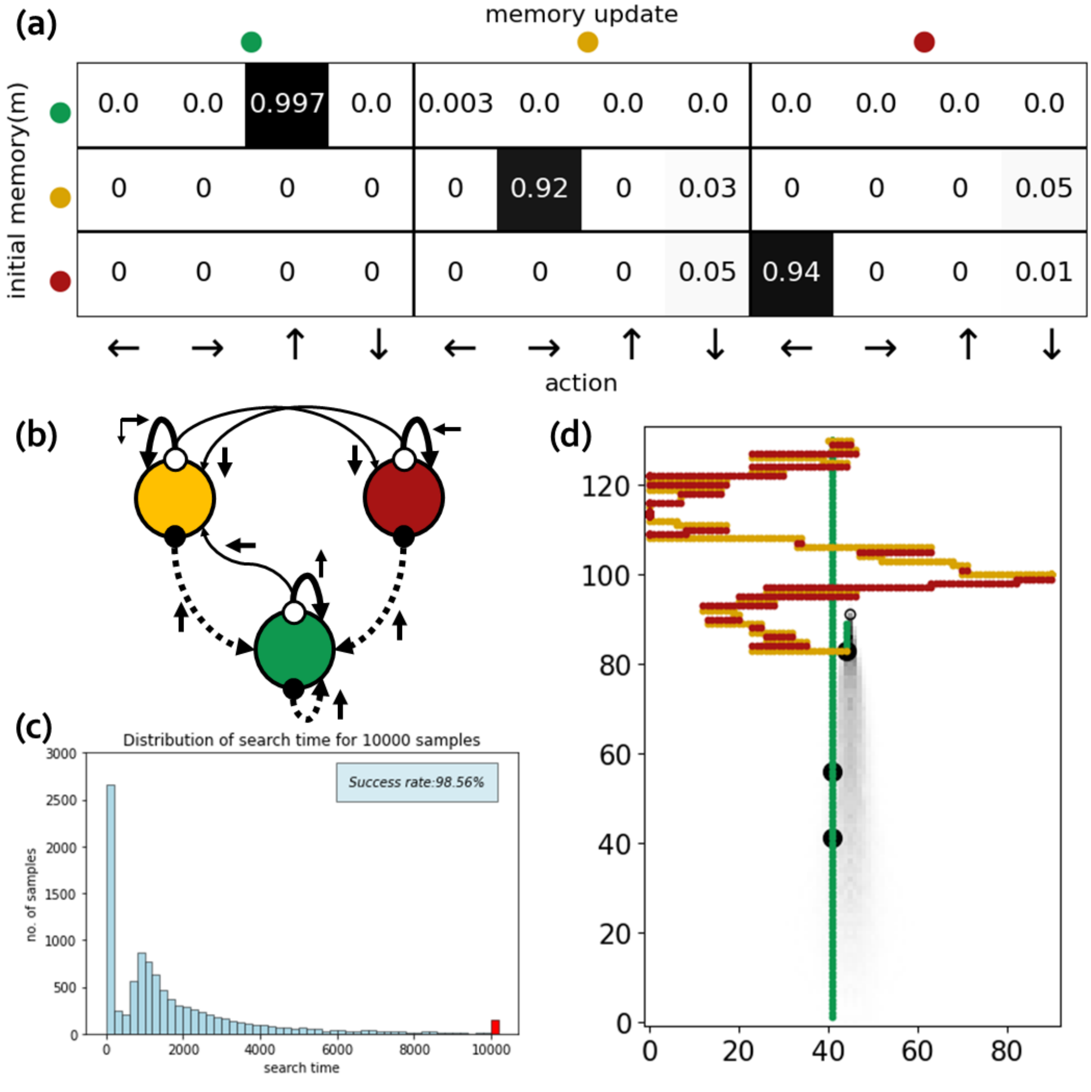


Figure 7: Finite state controllers with $|\mathcal{M}| = 3$ in a coarse grid with weak odor signal

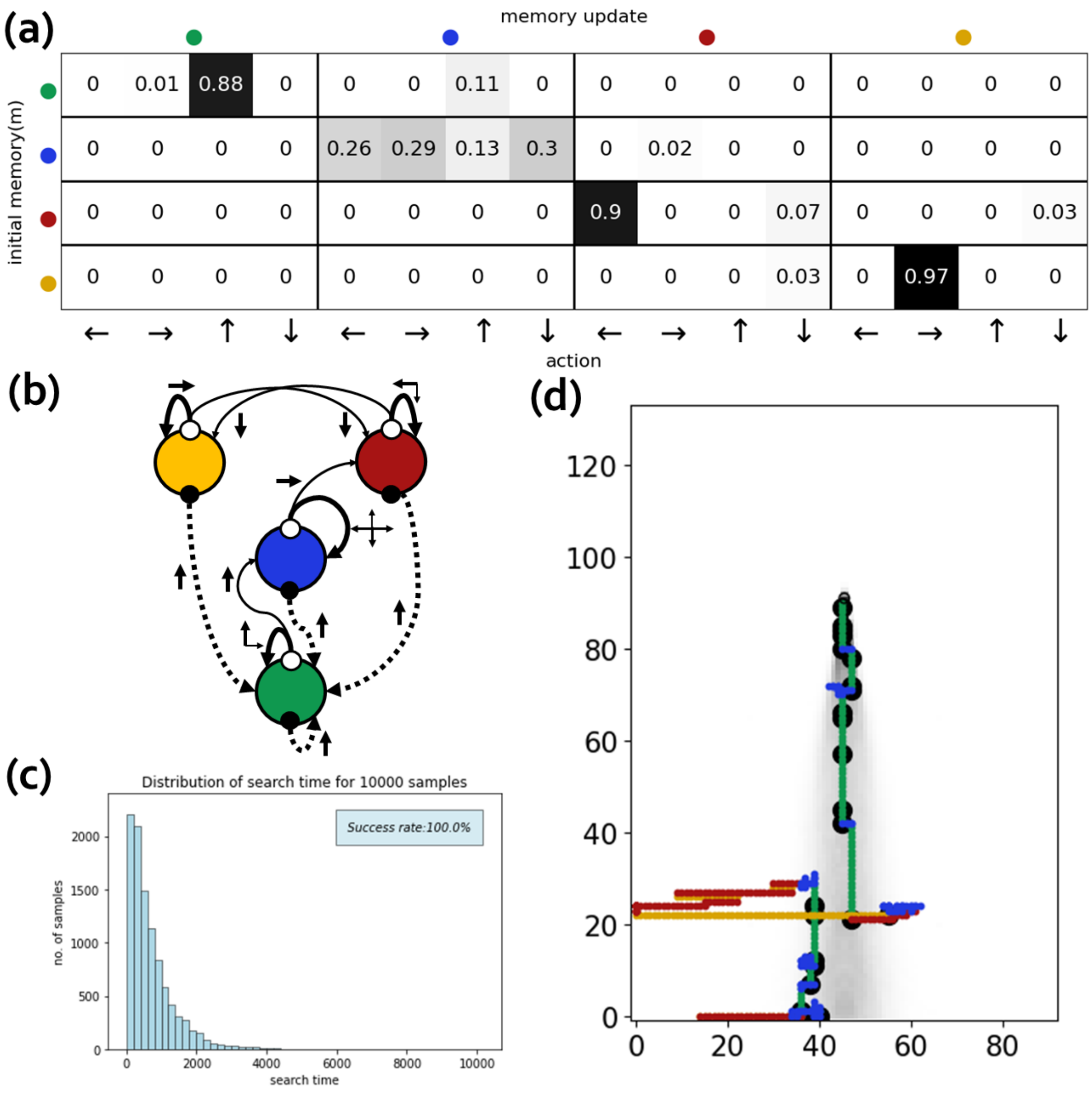


Figure 8: Finite state controllers with |$\mathcal{M}$| =4 in a fine grid with strong odor signal

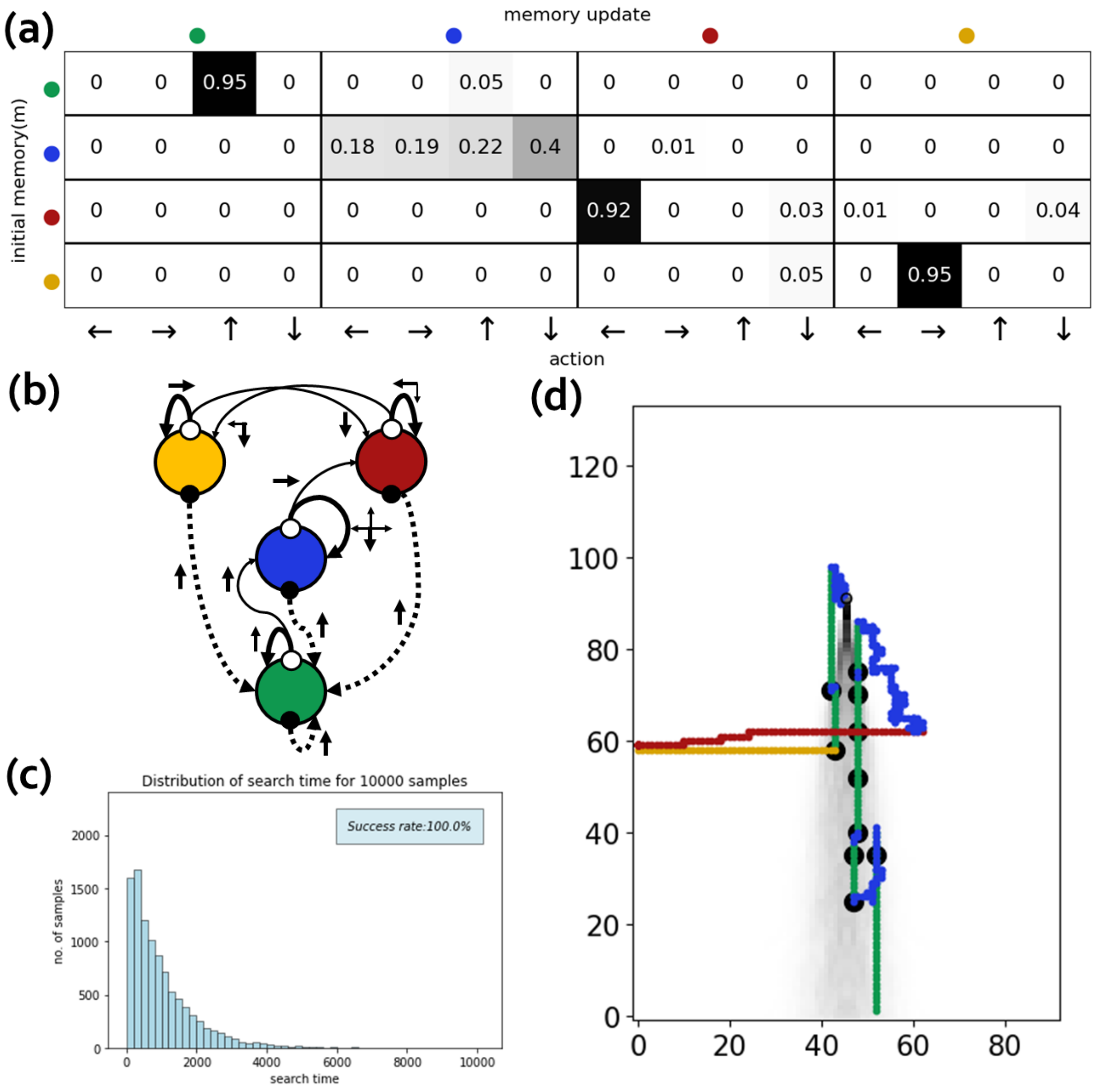


Figure 9: Finite state controllers with $|\mathcal{M}|$ =4 in a fine grid with weak odor signal

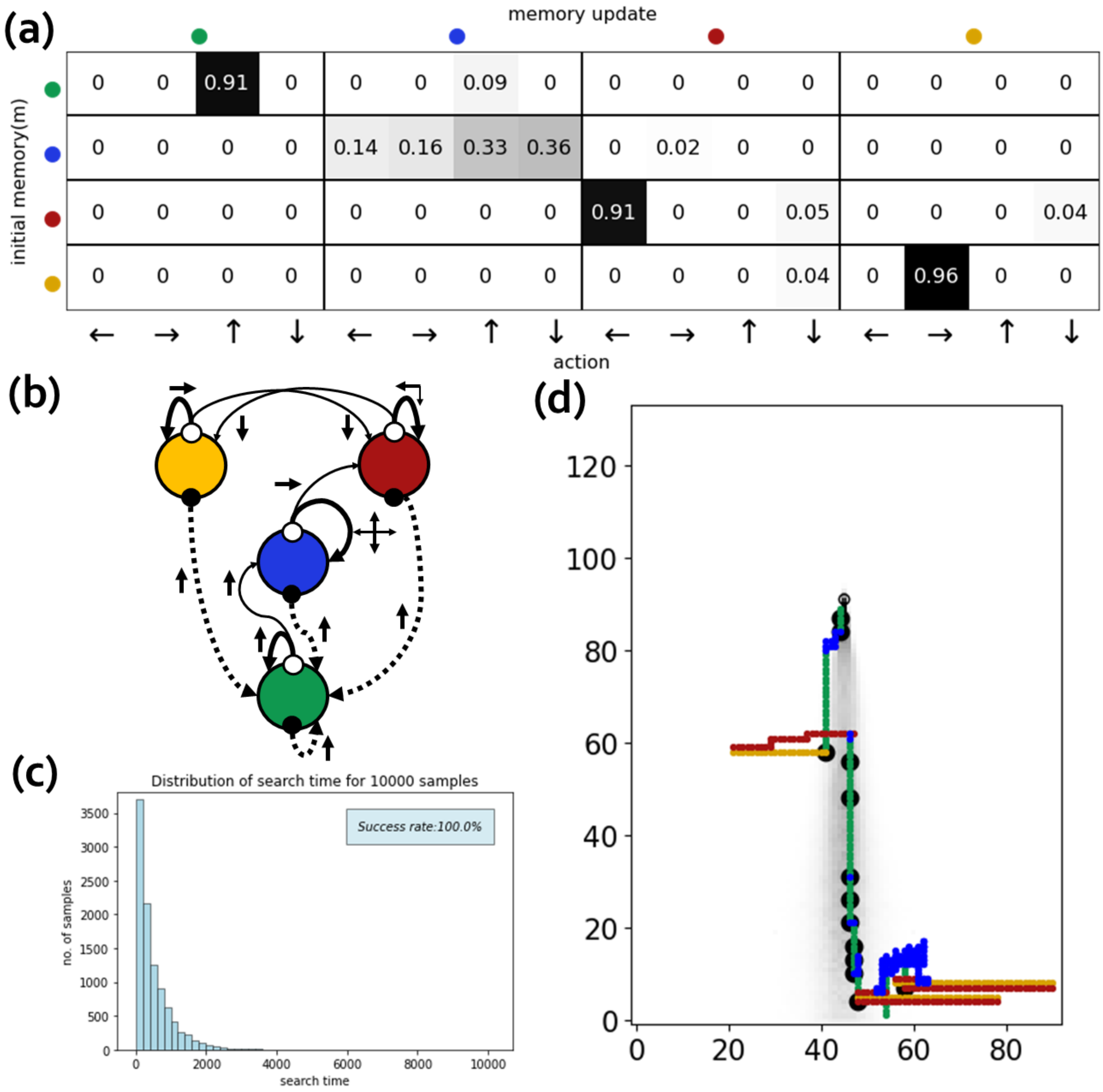


Figure 10: Finite state controllers with $|\mathcal{M}| = 4$ in a coarse grid with strong odor signal

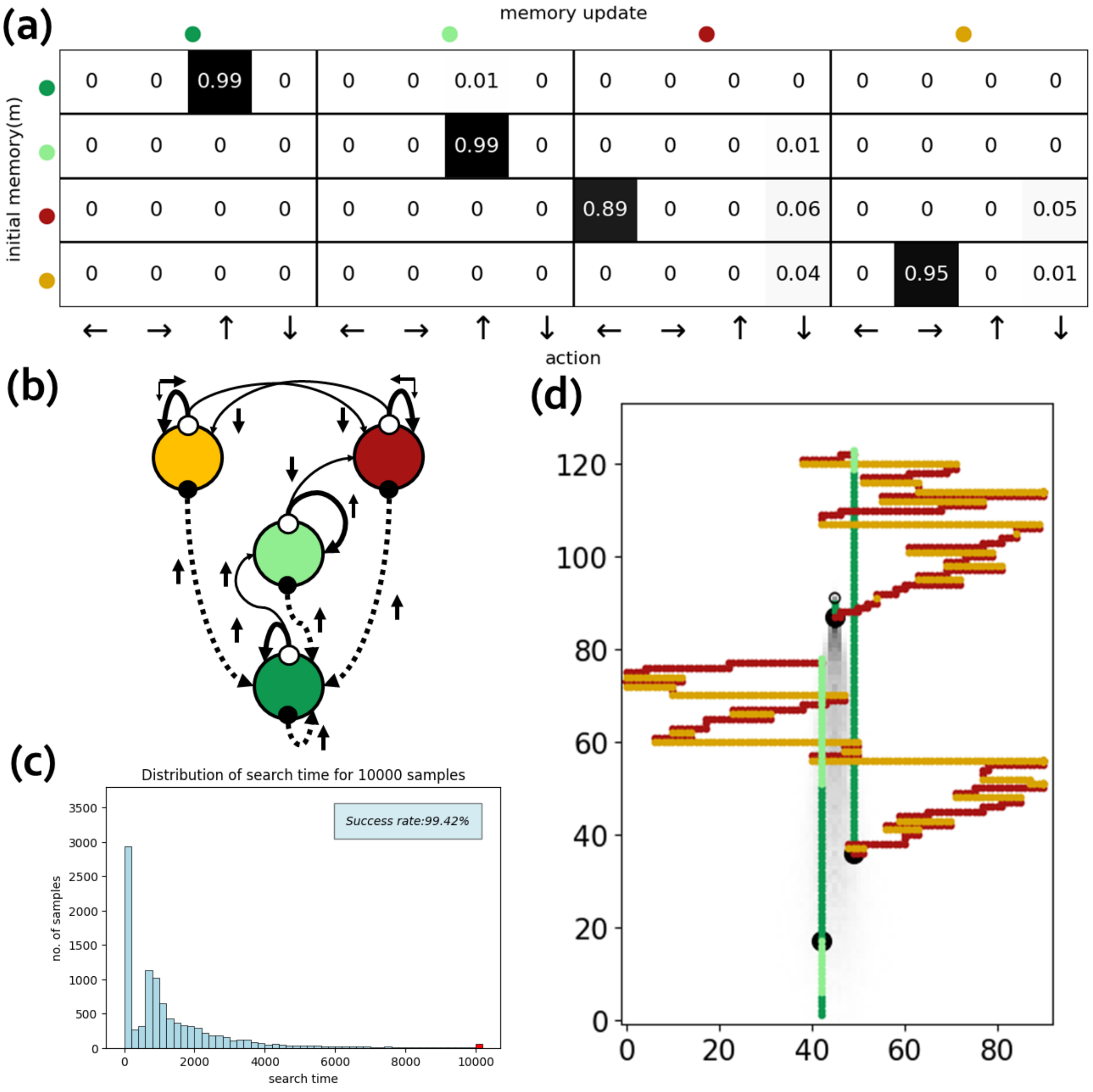


Figure 11: Finite state controllers with $|\mathcal{M}|$ =4 in a coarse grid with weak odor signal. Instead of the • state from the three previous controllers, we have now labeled the state as • to emphasize that it is associated to an upwind move, similarly to the • state.

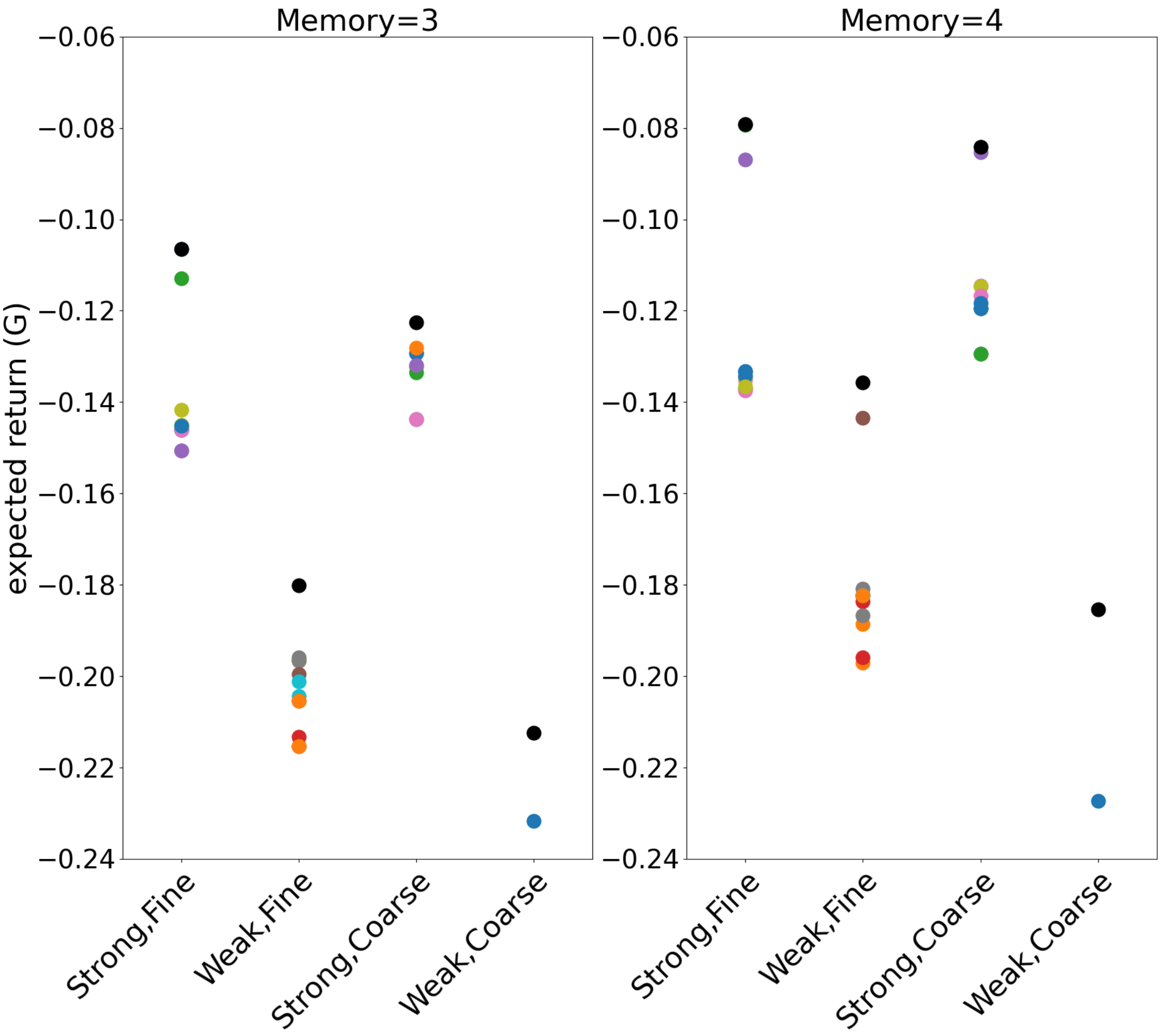


Figure 12: Performance after optimization for different initialization of parameters (shown as points of different color). The optimal policies presented in the main text and in Figures 4-11 are colored in black. We show here the expected return $G = r^T(1-\gamma T)^{-1}\rho$ for $\gamma = 0.99975$ and reward is equal to $-(1-\gamma)$ per time step – except for the case where the searcher is in the target location in which the reward is zero.
